# Comprehensive genome analyses of *Sellimonas intestinalis*, a potential biomarker of homeostasis gut recovery

**DOI:** 10.1101/2020.04.14.041921

**Authors:** Marina Muñoz, Enzo Guerrero-Araya, Catalina Cortés-Tapia, Ángela Plaza-Garrido, Trevor D. Lawley, Daniel Paredes-Sabja

**Affiliations:** Microbiota-Host Interactions and Clostridia Research Group, Departamento de Ciencias Biológicas, Facultad de Ciencias de la Vida, Universidad Andrés Bello, Santiago, Chile; Millennium Nucleus in the Biology of Intestinal Microbiota, Santiago, Chile; Host–Microbiota Interactions Laboratory, Wellcome Trust Sanger Institute, Wellcome Genome Campus, Hinxton, United Kingdom

**Keywords:** *Sellimonas intestinalis*, phylogenomic, gut homeostasis, extremely oxygen-sensitive species

## Abstract

*Sellimonas intestinalis* is a Gram positive and anaerobic bacterial species previously considered as uncultivable. Although little is known about this Lachnospiraceae family member, its increased abundance has been reported in patients who recovered intestinal homeostasis after dysbiosis events. In this context, the aim of this work was taken advantage of a culturomics protocol that allowed the recovery species extremely oxygen-sensitive from faecal samples, which led to the establishment of an S. intestinalis isolate. Whole genome sequencing and taxonomic allocation confirmation were the base to develop comparative analyses including 11 public genomes closely related. Phylogeographic analysis revealed the existence of three lineages (linage-I including isolates from Chile and France, linage-II from South Korea and Finland, and linage-III from China and one isolate from USA). Pangenome analysis on the established dataset revealed that although S. intestinalis seems to have a highly conserved genome (with 50.1% of its coding potential being part of the coregenome), some recombination signals were evidenced. The identification of cluster of orthologous groups revealed a high number of genes involved in metabolism, including amino acid and carbohydrate transport as well as energy production and conversion, which matches with the metabolic profile previously reported for healthy microbiota. Additionally, virulence factors and antimicrobial resistance genes were found (mainly in linage-III), which could favour their survival during antibiotic-induced dysbiosis. These findings provide the basis of knowledge about this species with potential as a bioindicator of intestinal homeostasis recovery and contribute to advance in the characterization of gut microbiota members with beneficial potential.

## Introduction

Gut microbiota plays important roles for human and other mammalian species, including: i) the maintenance of the structural integrity of the intestinal epithelial barrier;^1^ ii) the protection against the proliferation and colonization of enteropathogens;^2^iii) metabolites production or conversion of substances for the host;^3^ and iv) the stimulation of normal immune system functionality.^4^ All these functions are determined by diversity and abundance of microbial taxa that have been associated with host status (e.g. heath/disease, age, geographical origin among other comparison approaches). For this reason, the scientific community has been focusing its efforts on deciphering the composition of the microbial communities that inhabit this ecosystem.

Classical techniques to detect and study microorganisms involve *in vitro* culture, however, it is well known that most species inhabiting human gut cannot be cultured under standard conditions.^5^ To overcome this limitation, culture-independent DNA-based techniques, mainly based on next-generation sequencing (NGS), have been widely used to decipher almost all species at the intestinal level, that is the case of targeted NGS (tNGS) which has become the most popular scheme to depicting microbiota composition, thanks to the use of high-resolution markers to identify the taxonomic units (bacteria as well as eukaryotes and viruses), their variation among individuals or populations, and to infer phylogenetic relationships among the dominant taxa.^6^ This approach has been complemented with shotgun metagenomics technology, which also leads to describe microbiota composition, but in addition it allows to assembly whole genomes of the dominant taxa and to know the total content of nucleic acids present in the studied environment, which in the case of the gut, could provide informative markers of specific health/disease promoting factors.^7^

Studies based on culture independent NGS have shown that *Ruminococcaceae* and *Lachnospiraceae* are the most abundant Clostridial families at gastrointestinal tract of human and other mammals.^8, 9^ Although species diversity and the role of these two families are being studied, changes in their relative abundance have been observed in dysbiosis, being positively associated with healthy groups.^8^ In particular, the *Lachnospiraceae* family has gained interest during the last years due to the ecological adaptations exhibited by some of its species, associated with their capability to produce short-chain fatty acid (SCFA) during glucose fermentation.^10^ This capability attributed to commensal gut bacteria in healthy individuals^11^ has led to propose some *Lachnospiraceae* species as potentially beneficial gut microbiota member; however, few species of this family have been comprehensively studied.

One of the *Lachnospiraceae* species recently identified and poorly studied is *Sellimonas intestinalis*, a Gram positive and obligately anaerobic bacteria,^12^ initially considered as part of the gut microbiota fraction that remains uncultivated for its nature extremely oxygen-sensitive ‘EOS’.^13^ This limitation could have been the cause of the limited number of studies where *S. intestinalis* has been detected, being almost all aimed to deciphering the microbiome composition from a shotgun metagenomics approach.^13, 14^ In these studies, an increased relative abundance of *S. intestinalis* was detected in patients which recovered their intestinal homeostasis after suffering dysbiosis caused by chemotherapy treatment against colorectal cancer^15^ or therapeutic splenectomy of patients with liver cirrhosis.^16^ These findings suggest the potential of *S. intestinalis* as a biomarker candidate of gut homeostasis recovery. Conversely, punctual transversal studies have detected an incremented relative abundance of *S. intestinalis* in individuals with altered gut microbiota associated with chronic kidney disease^17^ and systemic-onset juvenile idiopathic arthritis.^18^ However, there are no studies aimed at clarifying the role of *S. intestinalis* within the intestinal microbiome.

A pivotal step to clarify the implication of *S. intestinalis* in host’s gut homeostasis is to know their genomic organization that allow to identify the genetic bases of their ecological role. However, due to *in vitro* culture limitations, only eight draft genomes have been obtained to November of 2019 (https://www.ncbi.nlm.nih.gov/genome/genomes/41970), that were assembled from shotgun metagenomics data. These genomes have been reported mostly from Eastern countries (China and South Korea), with a single genome reported from America (USA).^19^

For this reason, in this study we have employed a culturomics approach directed to isolate oxygen-sensitive intestinal microbiota species that allowed recovering *S. intestinalis*. Subsequently, a comprehensive whole genome analysis of this species was carried out to identify its genomic architecture, intra-taxa diversity, genetic population structure, potential metabolic profiles codifying for its genome and the presence of clinically important loci, as Virulence Factor markers (VFm) and antimicrobial resistance genes (AMRg), which could play a detrimental role in the colonization and relative abundance of this species in the complex intestinal environment. This approach represents an initial step to define the genomic bases that could support the role of this species in the intestinal microbiome and their potential as a biomarker of homeostasis gut recovery.

## Results

### Isolate establishment and biological source

A Gram-positive bacterial isolate with coccoid morphology (Supplementary Figure 1) was stablished under the conditions to recovery of microorganisms extremely sensitive to oxygen at gastrointestinal level standardized by our research group. The biological source of this isolate was a stool sample from a 23-year-old woman that despite being healthy at the time of sample collection, has a diagnosis of idiopathic rheumatoid arthritis, for this reason she was under treatment with Prednisone, a synthetic corticosteroid with glucocorticoid modulation, which support its anti-inflammatory effect and has proven as effective and safe for treatment of patients suffering this pathology.^20^ The individual consumes in addition *Chlorella* (a microalgae containing omega-3 fatty acids and carotenoids with antioxidant effect that have been proposed as a potential source of renewable nutrition),^21^ vitamin E with selenium and Korean ginseng. The individual did not use any antimicrobial treatment during the six months prior to the sample collection.

**Figure 1.**
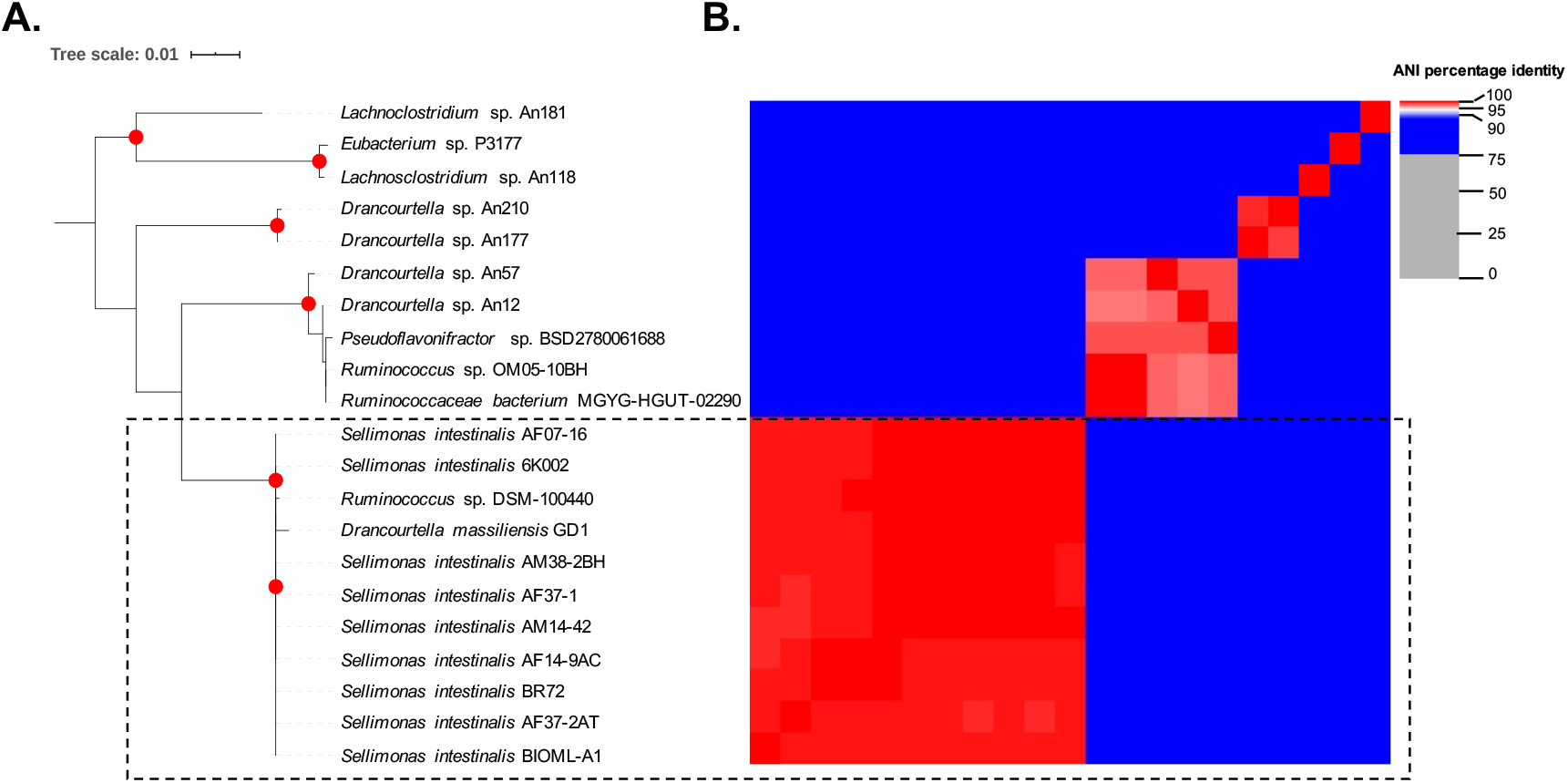
Taxonomic allocation analyses of studied genome using a phylogenomic approach. A) Phylogenetic reconstruction based on 16S-rRNA alignment for the 21 selected genomes. Sequences were aligned using MAFFT^55^ and then an approximately-maximum-likelihood phylogenetic tree was built in FastTree double precision version 2.1.10.^56^ Interactive Tree of Life V3 (http://itol.embl.de) was used for the graphic visualization.^61^ Red dots represent Bootstrap ≥ 90.0. B) Average Nucleotide Identity (ANI) analysis for the selected dataset. Two genomes with ANI results higher than 95% belong to the same microbial species. The analysis was developed using pyANI (https://github.com/widdowquinn/pyani).

### Assembly genome and taxonomic placement

The assembly genome obtained showed a length of 3,096,198 base pairs (bp), constituted by 32 contigs with an N50 of 3 and a length of 439,526 bp. The extraction and subsequent comparison of 16S-rRNA sequence revealed that the analyzed genome potentially belongs to one of the following genera: *Ruminococcus*, *Drancourtella* or *Sellimonas* (Supplementary Table 1). The search of reads of this species in ENA database, allowed to find the report for one isolate, that was assembly under the same conditions of the genome analyzed in this study. The analysis of 2,902 genomes of *Ruminococcaceae* and *Lachnospiraceae* genomes used for the revision of *Clostridiales* order study allowed to identify which the analyzed genome and the ENA report makes part of a node well supported which included other 9 genomes of *Sellimonas intestinalis* (Supplementary Figure 2). These 11 genomes were then considered as the *S. intestinalis* node. Interestingly, two incongruencies in taxonomic allocation of publicly available genomes were detected, being previously deposited as *Ruminococcus* sp. DSM-100440 and *Drancourtella massiliensis* GD1, and consistently clustered with the genome set under study (16S-rRNA phylogenetic reconstruction (Fig. 1A) and ANI analysis (Fig. 1B)), that hereafter will be treated as part of *S. intestinalis* node. This well supported node was pruned join the closet node (with 7 genomes), that included mostly *Drancourtella* genomes, and for that was identified as *Drancourtella* node. Within this node were also found incongruences in taxonomic allocation, being included two *Ruminococcus* and one *Pseudoflavonifractor* genomes (Supplementary Figure 2). Three additional representative genomes clustering in related nodes were included as outgroups (*Lachnosclostridium* sp. An181, *Eubacterium* sp. P3177 and *Lachnosclostridium* sp. An118). Under these parameters a set of 21 assemblies were included in the data set for subsequent analysis.

**Figure 2.**
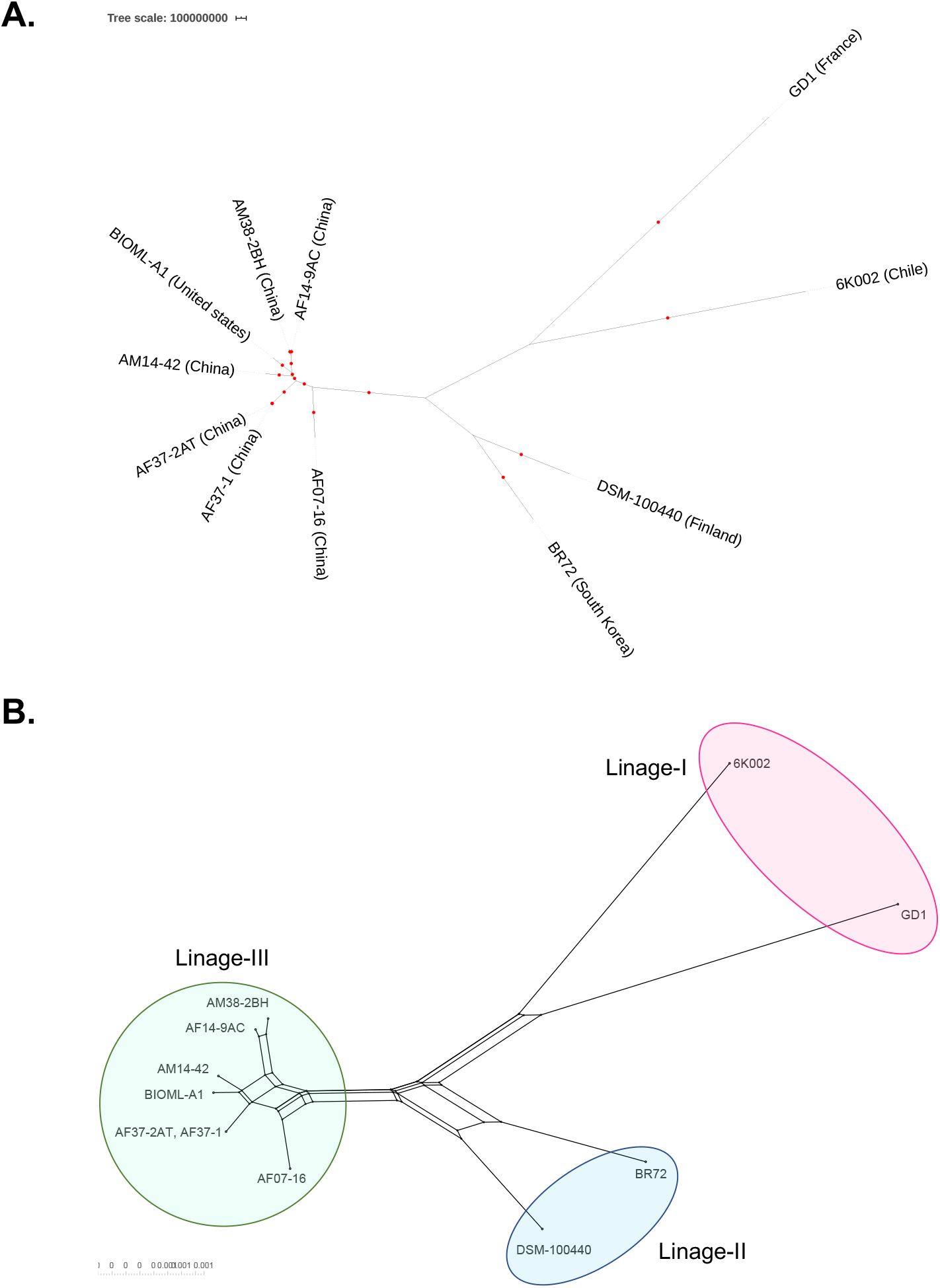
Phylogeographic analysis and phylogenetic networks used to predict the genetic population structure of *Sellimonas intestinalis*. A) Bayesian evolutionary analysis based on Markov Chain Monte Carlo (MCMC) implemented in BEAST-2^60^ carried out from the core genome alignment (with 22,453 positions of length) of the 11 sequences selected assemblies. GTR substitution model was chosen as the best model in jModelTest v0.1.1.^58^ B) phylogenetic network using neighbor-net method conducted in SplitsTree5.^62^

The phylogenetic reconstruction based on 16S-rRNA alignment for the 21 selected genomes showed that the 11 genomes previously assigned to *S. intestinalis* node remain clustered together (Fig. 1A). These findings were compared with the ANI percentage identity which was higher than 95% for all these 11 *S. intestinalis* genomes (Fig. 1B), which led to verify that under the traditional criteria to identify microbial species from whole genome data (16S-rRNA and ANI), all 11 assemblies correspond to *S. instestinalis* (Fig. 1A and 1B). The information on the genomes included in *S. intestinalis* is described in the Supplementary Table 2.

### Intra-species diversity and genetic population structure

A preliminary BLAST comparison of 11 *S. intestinalis* selected assemblies revealed a high level of genome conservation; however, some genome regions were differentially present in groups of isolates. The map comparing the complete genomes delimited by these populations is described in the Supplementary Figure 3. As next step, the pangenome analysis of *S. instestinalis* dataset showed a codifying potential of 4,627 genes (Supplementary Table 3), which are almost equally distributed between core genes (*n*=2,318; 50.1%) and accessory genes (*n*=2,309; 49.9%).

**Figure 3.**
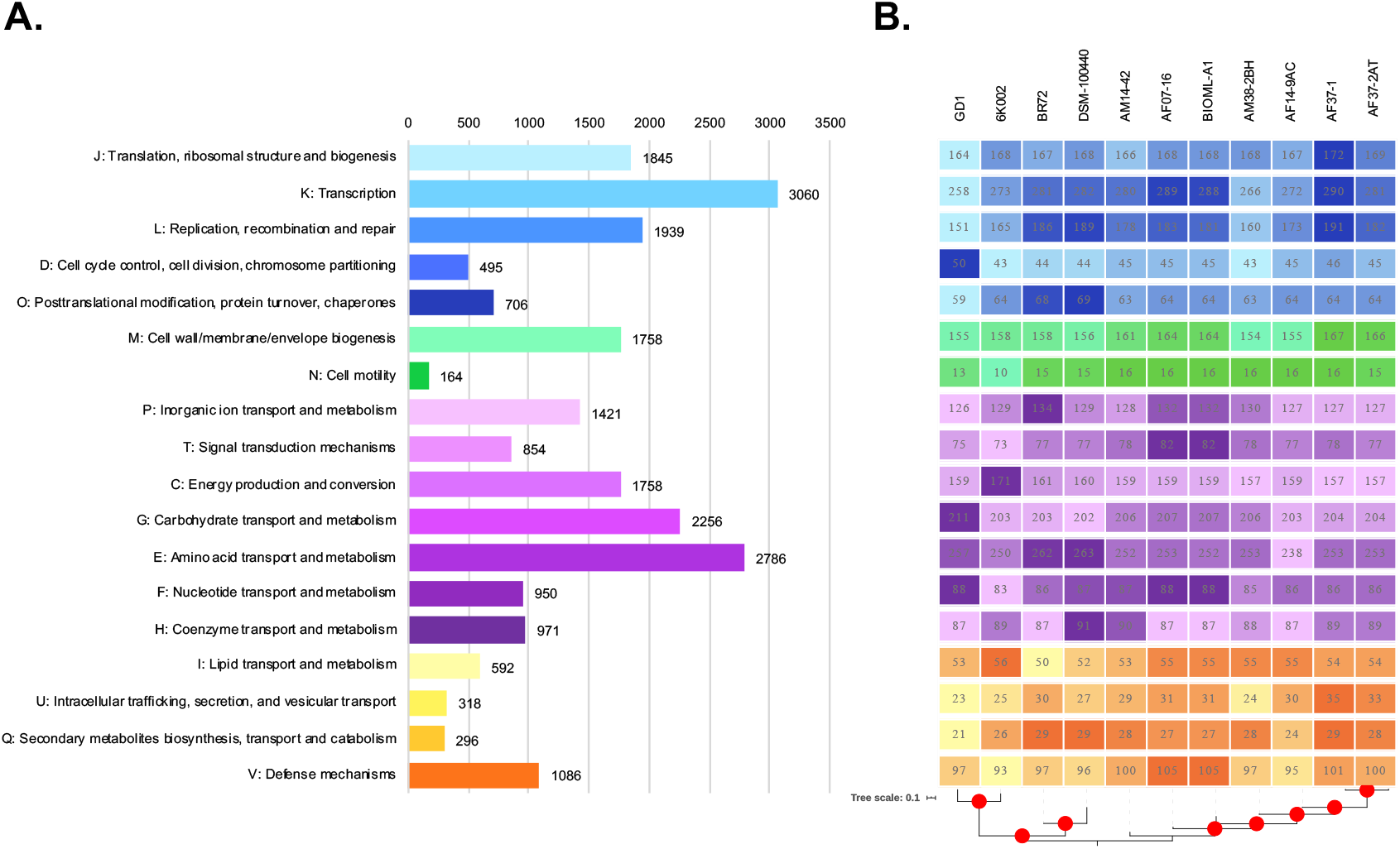
Cluster of orthologous groups (COGs) for A) global data set and B) individual isolates. eggNOG-mapper v2 was used as a tool for fast functional annotations of sequence collections.^63^

A phylogeographic analysis, based on a Bayesian evolutionary approach, was conducted from coregenome alignment (with 22,453 positions of length) of the 11 sequences selected assemblies. The phylogenetic three topology revealed that *S. intestinalis* could have diversified into at least three major linages with a possible relation by geographical origin (Fig. 2A). The first linage (Linage-I) included isolates from Chile and France, while the second linage (Linage-II) isolates from South Korea and Finland, and the third linage (Linage-III) isolates majority from China and only one from the USA. This population genetic structure was confirmed by the phylogenetic network topology that showed that although there are recombination signatures (indicated by reticulation events observed), the three linages detected by phylogeographic analysis are divergent among their, which supports the hypothesis of the existence of three main populations within this species (Fig. 2B).

### Cluster of orthologous groups

To explore the coding potential of the genome set under analysis, firstly a COGs was developed for both global data set (Fig. 3A) and individual isolates according to the linages to them belong (Fig. 3B). The results showed that this species directs much of the coding potential to essential biological processes such as transcription, translation and replication. However, it can be observed that an important part of their genes could be involved in metabolism, including amino acid and carbohydrate transport as well as energy production and conversion (Fig. 3A). Differential profiles were detected in the identified populations, finding that the linage-I and linage-II clusters have more genes involved in metabolic processes, while linage-III isolates revealed profiles with more genes involved in the cell cycle, intracellular trafficking, secretion, and vesicular transport (Fig. 3B).

### Virulence factors and antimicrobial resistance genes

Considering that about half of the genes coding for this species are part of the accessory genome, we inspect the genes differentially transported by the lineages detected (Fig. 4A). This analysis allowed to identify that the clustering in three populations is maintained in the phylogenetic reconstruction based on the accessory genome, as was found in coregenome phylogenetic analysis (Fig. 2A).

**Figure 4.**
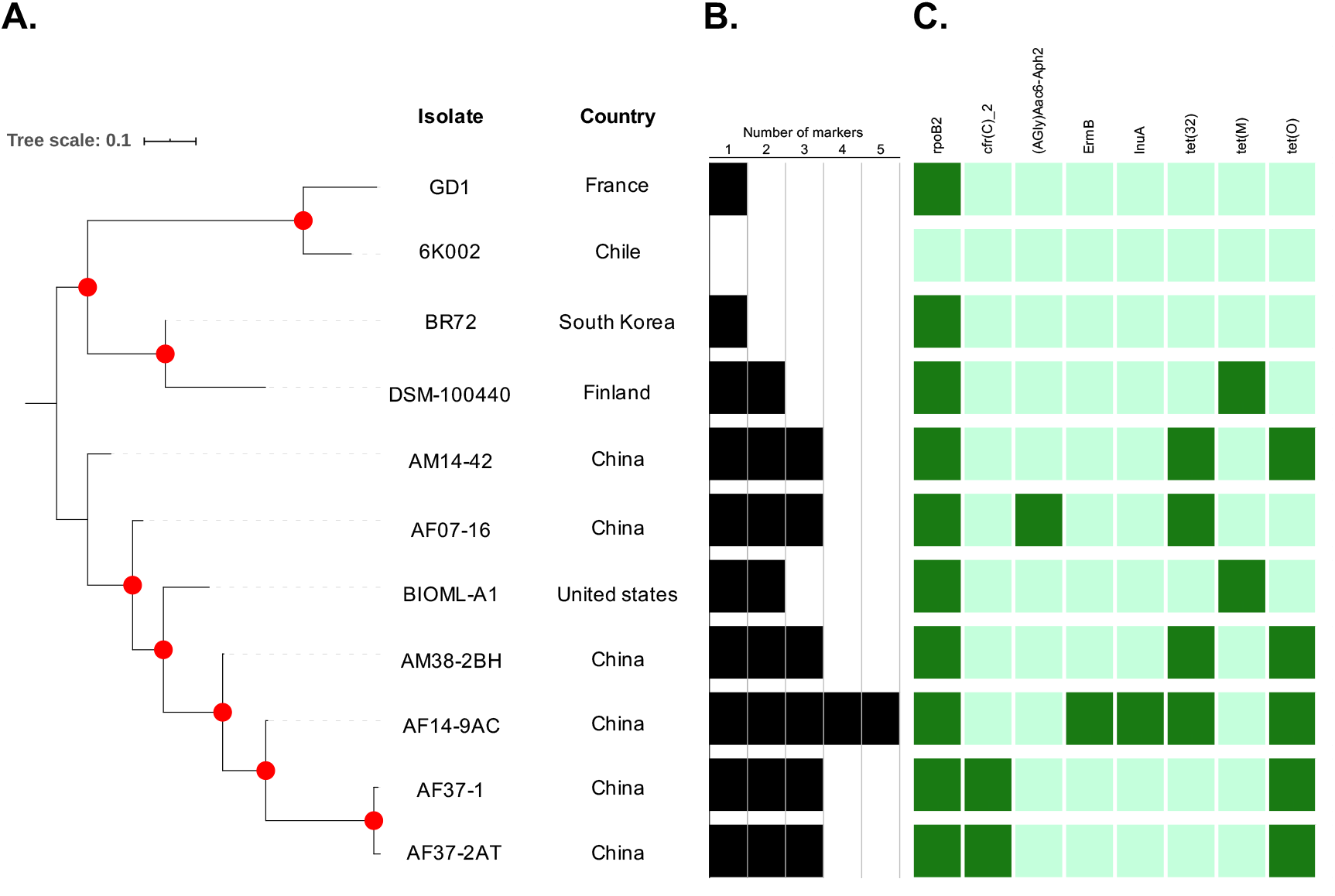
Virulence factors and antimicrobial resistance genes detected in *Sellimonas intestinalis* genomes. A) phylogenetic reconstruction from accessory genome alignment. B) frequency of markers found in each assembly; and C) presence-absence matrix which describe the markers detected in each genome. Abricate 0.8.4 (https://github.com/tseemann/abricate) was used to make BLAST against the sequences previously reported in the following databases: CARD,^64^ Resfinder,^65^ NCBI,^66^ ARG-ANNOT,^67^ VFDB^68^ and PlasmidFinder.^69^

VFm and AMRg are important loci for survival of bacterial species because could modulate changes in their abundance under different biological contexts and the subsequent transmission dynamics between hosts (Fig. 4B). Although the exhaustive search from both assemblies and reads revealed that the isolate from Chile (linage-I) not carry known VFm nor AMRg, the extended search of this loci from assemblies included in the comparative dataset reveled that the other genome clustering in the same linage-I from France carry *rpoB2* marker, associated with resistance to rifampin resistance. *rpoB2* was found in all other 9 evaluated genomes. *tet*(M) marker (associated with tetracycline resistance) was the only one additional marker found in linage-II, being transported by the isolate from Finland. Interestingly, linage-III exhibited the greatest amount of AMRg, being found from 2 to 5 (in the case of AF14-9AC from China) genes per genome. Among the genes with higher frequency were found: *tet* elements (*tet*32 and *tet*(O)), present in five and four genomes, respectively, and *cfr(C)_2* (conferring linezolid resistance) present in two genomes. In addition, (AGly)Aac6-Aph2, associated with aminoglycoside drug class resistance, *ermB* conferring macrolide-lincosamide-streptogramin antibiotic resistance, and *lnuA* associated to lincosamide resistance, all of these present in a single genome each one.

## Discussion

Recent studies based on amplicon-based sequencing and shotgun metagenomics have contributed to the description of diversity and abundance of gut microbial communities ^22^ and it has even been possible to propose associations with host states ^23^ and make inferences about the possible functions of specific members of this complex ecological network.^24^ However, genomic characterization of gut microbiota members represents a challenge to deciphering the genetic bases supporting the biological function of microbial species inhabiting gut, being an essential initial step their recover by *in vitro* culture, that have an increased complexity for EOS species.^25^ For this reason, this work describes the isolation and genomic features of *S. intestinalis*, a understudied *Lachnospiraceae* species recovered during a culturomics approach directed to recover EOS species within microbiome environment.

During a genomic characterization study is essential a precise taxonomic allocation of target genomes and those included in the comparative dataset to avoid mistakes in the biological inferences. In this study, inconsistencies in taxonomic classification were detected at different levels: i) in the allocation of species to families with little phylogenetic relationship, as is the case of *Clostridium difficile* that had been included within the *Clostridiaceae* family, but after detailed analysis of the Phylogenetic relationships were classified within the *Peptostreptococcaceae* family,^26^ or ii) in the taxonomic assignment of individuals, as revealed even before this work for *S. intestinalis*, which in other works had previously been detected as *Ruminococcus*, but later of the sequencing of its complete genome, it was correctly assigned.^13^ These types of findings reveal limitations in the traditional analysis schemes of complete genome data and make clear the need for further studies that lead to clarify the classification of under-studied anaerobic families.

The study of genetic population structure represents an important tool to determine the population sizes, dispersal potential and evolutionary rates lead over geographical scales during characterization of a microbial species.^27^ For the case of *S. intestinalis*, the limited number of isolates analyzed represents a limitation, for example, some remarkable profiles were identified, as is the case of the Sin-II and Sin-III lineages in which the isolates that comprise them show a slight degree of divergence (Fig. 2A and 2B), so that the increased number of individuals could lead to the identification of independent populations. These findings were supported by pangenome results that revealed that despite the limited number of genes of core genome (*n*=2,318, 50.1%), this could be a first indicator of the high intra-taxa diversity of this species. This type of finding has been detected in species such as *Pseudomonas aeruginosa*,^28^ a species of interest in health that exhibits a high frequency of gene loss and gain. The pangenome data also allowed the evaluation of phylogenetic relationships from the coregenome (Fig. 2A), which led to the detection of three potential linages, which were subsequently ratified through the construction of phylogenetic networks (Fig. 2B), which showed that despite the potential recombination events, supported by crosslinking in the networks, these populations could be highly divergent among them. Interestingly, possible common geographical origins were identified among the observed populations, with the exception of the USA isolate that clustering with isolates from China (Sin-III linage), that could be attributed to migration of human population, as has been identified for other pathogens.^29^

The effect of specific members over gut microbiome composition have been attributed mainly to their metabolic profiling in which some subproducts can stimulate specific process in the complex gut environment.^30^ For that reason, the metabolic profiling of *S. intestinalis* from whole genome data using COGs analysis was determinant to deciphering that the genes required for the survival of the bacteria (Fig. 3A), the genes associated with amino acid and carbohydrates transport and metabolism were highly frequent in this species. The individual analyzes grouped by the detected populations, showed that the isolates of oriental origin (linage-III), codify a greater number of genes of AMRg than the other geographical origins. These results are of importance, because these types of genes are determinants for the realization of metabolic processes that lead to the production of short-chain fatty acid (SCFA) during glucose fermentation,^10^ mainly butyric acid,^31^ which It has been proposed as one of the characteristics of the candidate members for healthy microbiota.^11^ In the specific case of the patient from whom the isolate studied in this work was obtained despite the immunosuppression caused as a possible consequence of the anti-inflamatory effect of Prednisone (the corticosteroid consumed as treatment of rheumatoid arthritis)^20^, the healthy lifestyle habits and the consumption of substances with potential restorative effect of the gut microbiota as Chlorella (the microalgae consumed by the donor individual because has shown potential as antioxidant and treatment for different health conditions)^21^, they could contribute to the restored effect of the Microbiota, and the presence of *S. intestinallis* could then be complying with the hypothesis as a biomarker of the recovery of intestinal homeostasis.

The isolation and characterization of microbiota members contribute to deciphering the genome bases of their effect in the gut microbial ecology,^32^ as well as to detect members that potentially play a role as reservoirs of antibiotic resistance.^33^ In the particular case of *S. intestinalis*, several genes associated with antibiotic resistance, such as *rpoB2* in most of the analyzed isolates, and mobile genetic elements, such as the family of *tet* AMRg (Fig. 4). These findings could represent the basis for the survival of this species at the intestinal level, despite the adverse conditions that this niche naturally represents. Additionally, it could explain its role as a biomarker, after the presentation and restoration of homeostasis after dysbiosis generated by different causes. The relevance of these findings should be subsequently confirmed through the application of phenotypic tests that lead to identify the minimum inhibitory concentrations at which the proliferation of this species is inhibited *in vitro*.

Considering the differential presence of genes biological and clinically relevant among the three *S. intestinalis* linages found in this work, is necessary to develop future studies conducting to develop a molecular typing method to quickly identify isolates and which contributes to clarifying the phylogenetic relationships and evolutionary history of this species. Additionally, taking into account that this study was aimed at analyzing data from complete genomes of *S. intestinalis* and no phenotypic tests were performed, it is necessary to carry out further studies that lead to identify the impact of the expression of these VFm and AMRg and their potential role in modulation of the relative abundance of this species under different biotic contexts. Despite this limitation, the identification of these markers could support the hypothesis that some members of the microbiota could fulfill resistance reservoir-function from which bacterial pathogens can acquire resistance is the human gut microbiota,^13^ generating interest at the health level. This approach represents the first step conducing to the genomic bases that support *S. intestinalis* survival under conditions of dysbiosis and subsequent proliferation after the homeostasis reestablishment that could play an important role in maintaining the optimal conditions for host development.

## Materials and Methods

### Sample collection

A culturomics approach was applied to stool samples from adult Chilean individuals, within the framework of the project Millennium Nucleus in the Biology of Intestinal Microbiota, aimed at detecting and characterizing the microorganisms that make up the intestinal microbiota of healthy individuals in Latin America, developed by our research team. This project was approved by Comité de Bioética de la Facultad de Ciencias de Vida, Universidad Andrés Bello, through the act 013-2017. All patients enrolled in this study agreed to participate and signed an informed consent form.

### Bacterial isolate recovery

This approach involved the optimization of a protocol for Extremely Oxygen-Sensitive (EOS) intestinal bacteria isolation as follow: stool samples (collected in sterile containers without preservation media) were processed within the first 72 hours. Next, the samples were mechanically homogenized and divided into two fractions that were treated independently. The first (approximately 50%), was washed with 100% ethanol to reach 70% (w/v) and incubated for 4 hours in anaerobiosis. The biological material was then precipitated by centrifugation, to discard the ethanol, and then washed twice with sterile molecular grade water. The second fraction of the sample was processed without washing. The two fractions were weighed and then independently resuspended in sterile 1X PBS (1mL per 100mg of feces), to be then serially diluted (from 10^−1^ to 10^−5^ for the sample washed with ethanol and from 10^−1^ to 10^−8^ for the sample processed directly). Each dilution of the two treatments was seeded in duplicate on the complex and broad-range YCFA medium,^34^ in two formats: traditional or supplemented with taurocholate (Winckler) (0.1% v/v). Finally, they were incubated for 72-96 hours at 37 °C under anaerobic conditions. The manipulation and incubation of samples were conducted in an anaerobic chamber Bactron EZ2 (ShellLab).

The colony forming units (CFU) obtained were streaked on YCFA plates, and after 24-48 hours of incubation under the conditions described, and then their quality and morphology were evaluated by classical microbiological techniques (macro and microscopic observation). The verified colonies were propagated in liquid YCFA medium to increase their biomass to establish the isolates, using the same incubation conditions. This isolate was named 6K002.

### DNA extraction and whole genome sequencing (WGS)

The biomass recovered from isolate incubation in broth medium was subjected to DNA extraction using the commercial kit Wizard® Genomic DNA Purification Kit (Promega Corporation, Madison, WI, USA), following the manufacturer’s recommendations. The DNA sequencing was carried out by Wellcome Trust Sanger Institute on an Illumina HiSeq 2000 platform, with a read length of 100 bp, under the “developing and implementing an institute-wide data sharing policy following conditions”.^35^

### Genome assembly and quality control verification

The reads obtained from WGS were *De novo* assembled using Unicycler v0.4.8, an assembly pipeline for bacterial genomes defined as a SPAdes-optimiser (Spades v3.13.1) which generates the best possible assembly,^36^ using parameters by default. The quality of the genome assembly was evaluated using the GenomeQC_Filter_v1-5 script, ^37^ which considers as parameters the maximum number of contigs per genome (fixed to 400) and a maximum size of each genome (considering 8 Mbp) and then extracts the small subunit 16S rRNA gene sequences (16S-rRNA).

### Taxonomic placement and data retrieval

Initially, the 16S-rRNA sequence previously extracted was used to search sequence similarity against the data available in public datasets using the BLASTn algorithm [39], results that were subsequently verified by 16S-rRNA sequence alignment using SILVA Incremental Aligner (SINA) service.^38^

Next, a dataset with 2,902 *Ruminococcaceae* and *Lachnospiraceae* genome assemblies, publicly available in PATRIC,^39, 40^ ENA^41^ and NCBI^42^ databases and which passed the assembly quality test previously described, were analyzed to identify the genomes most related with the analyzed assembly. This dataset makes part of a parallel work of our research team directed to evaluate the phylogenetic relationships of Clostridiales order. Parallelly, a search of reads for ‘*Sellimonas’* genus was conducted in the European Nucleotide Archive (https://www.ebi.ac.uk/ena/data/search?query=Sellimonas), with the aim of recovering the greatest number of genomes for analysis. The obtained reads were subject to the genome assembly and quality control verification methodology describe in the previous section.

The complete genome dataset was the base to select the node closely related to the analyzed genomes, throughout phylogenetic reconstruction based on 16S-rRNA sequence under the parameters described in the corresponded section. The set of assemblies selected were subjected to a step of delimiting species using average identity of nucleotides (ANI),^43^ using pyANI 0.2.10, a Python3 module and script that provides support for calculating average nucleotide identity (ANI) and related measures for whole genome comparisons, and rendering relevant graphical summary output (https://github.com/widdowquinn/pyani).^44^ pyANI analyses was developed using blast and other settings by default. Scores of ANI higher than 95.0%, were used to verify that the genomes belong to the same species.

A graphical map of the genome assemblies identified as belonging to the same species of studied genome, was built in the CGview server,^45^ where a comparison was made in pairs to identify the differences between the genomes, using the tool based on the BLAST algorithm, included inside the server.

### Annotation and pangenome analysis

An automated annotation pipeline was applied to the complete set of evaluated genomes. This pipeline is based on Prokka v1.13,^46^ as follows: Infernal v1.1.2^47^ was run to predict RNA structures, followed by an analysis in Prodigal v2.6.3^48^ to predict proteins. Aragorn v1.2.38^49^ was used to predict tRNAs and tmRNAs, and Rnammer^50^ was used to predict ribosomal RNAs. All predicted genes were then annotated throughout databases search following this order: genus specific databases were generated by retrieving the annotation from RefSeq.^51^ The protein sequences were then merged using CD-hit version 4.8.1^52^ to produce a non-redundant blast protein database. Next UniprotKB/SwissProt^53^ was searched, considering kingdom specific databases for Bacteria. The complete set of genomes evaluated was submitted to the aforementioned annotation pipeline.

As a next step, the pangenome was determined using the Roary tool version 3.11.2,^54^ taking as definition of coregenome a percentage identity of 95% using Protein-Protein BLAST 2.9.0+ and the presence in 99% of the analyzed genomes.

### Phylogeographic analyses

The phylogenetic relationships among *Ruminococcaceae* and *Lachnospiraceae* assemblies was evaluated to identify the data most closely related with the studied genome. For that, the 16S-rRNA sequences extracted during the quality control verification step were aligned using MAFFT v7.407^55^ using parameters by default and then an approximately-maximum-likelihood phylogenetic tree was built in FastTree double precision version 2.1.10^56^ with settings by default. The robustness of the nodes was evaluated using the Bootstrap method (BT, with 1,000 replicates).

After the definition of dataset to analyze, the phylogenetic relationships among isolates were evaluated using a Bayesian evolutionary approach based on Markov Chain Monte Carlo (MCMC) implemented in Beast v1.10.4^57^ from the pangenome alignment (with a length of 22,453 nucleotides) of the 11 sequences selected assemblies. GTR substitution model was chosen as the best model in jModelTest v0.1.1,^58^ an uncorrelated relaxed clock model and the skyline population model, were considered initial parameters. Twenty independent MCMC was carried out, each with a chain length of 100,000,000 states and resampling every 10,000 states. Log files were summarized with Tree Annotator v2.4.8 [44] using 10% burning. The effective sample size (ESS) were >200 for all parameters; convergence and mixing were assessed using trace plot in Tracer v1.7.1.^59^ Tree files generated were summarized with Tree Annotator v2.4.8^60^ using 10% burning, with maximum clade credibility and node heights at the heights of common ancestors. A node dating step was conducted using isolate metadata (date of isolate and geographic origin). The graphic visualization of all phylogenetic trees was obtained in the web tool Interactive Tree of Life V3 (http://itol.embl.de).^61^ Additionally, phylogenetic networks were conducted with the aim to detect recombination signatures in the analyzed population. This analyses were carried out in SplitsTree5,^62^ using neighbor-net method.

### Codifying potential of *S. intestinalis* genome

The annotation outputs were additionally used to identify Clusters of Orthologous Groups (COG) using eggNOG-mapper v2 under default settings, a tool for fast functional annotations of sequence collections.^63^ The COG categories were subsequently were represented in a histogram.

Virulence factor markers (VFm) and antimicrobial resistance genes (AMRg) were identified from whole genome assembles using Abricate 0.8.4 (https://github.com/tseemann/abricate), making BLAST against the sequences previously reported in the following databases: CARD (1,749 sequences, last update: Jul 8, 2017),^64^ Resfinder (1,749 sequences, last update: Jul 8, 2017),^65^ NCBI (1,749 sequences, last update: Jul 8, 2017),^66^ ARG-ANNOT (1,749 sequences, last update: Jul 8, 2017),^67^ VFDB (1,749 sequences, last update: Jul 8, 2017)^68^ and PlasmidFinder (1,749 sequences, last update: Jul 8, 2017)^69^. Minimum DNA identity of 75% was used as detection thresholds. As a confirmation step about VFm and AMR presence, Ariba (Antimicrobial Resistance Identification By Assembly) version 2.0^70^ was run from reads of the studied isolate.

## Acknowledgements

The authors thank Wellcome Trust Sanger Institute, in particular to the core library and sequencing teams for whole genome sequencing of *Sellimonas intestinalis* GK002 and pathogen informatics team for the use of several automated pipelines during the processing and analysis of the whole genome sequence data.

## Supplementary material legends

**Supplementary Figure 1.** Microscopic morphology of *Sellimonas intestinalis* isolate. The results indicate a Gram-positive bacterial with coccoid morphology.

**Supplementary Figure 2.** Phylogenetic reconstruction of 2,902 genomes of *Lachnospiraceae* and *Ruminococcacea* genomes based on 16S-rRNA alignment which allowed to define a node well supported which included the studied assembly and other 9 genomes.

**Supplementary Figure 3.** Graphical map of the 11 genome assemblies identified as belonging to the same species of studied genome, built in the CGview server.^45^

**Supplementary Table 1.** Comparison of 16S-rRNA sequence using BLAST which revealed that the analyzed genome belongs to one of the following genera: *Ruminococcus*, *Drancourtella* or *Sellimonas*.

**Supplementary Table 2.** Information of *S. intestinalis* genomes included in comparative analyses.

**Supplementary Table 3.** Pangenome analysis of *S. instestinalis* dataset showed a codifying potential of 4,627 genes.

