## Supplementary figures and images for "Comprehensive genome analyses of *Sellimonas intestinalis*, a potential biomarker of homeostasis gut recovery"

### Sup_Fig_1.tiff

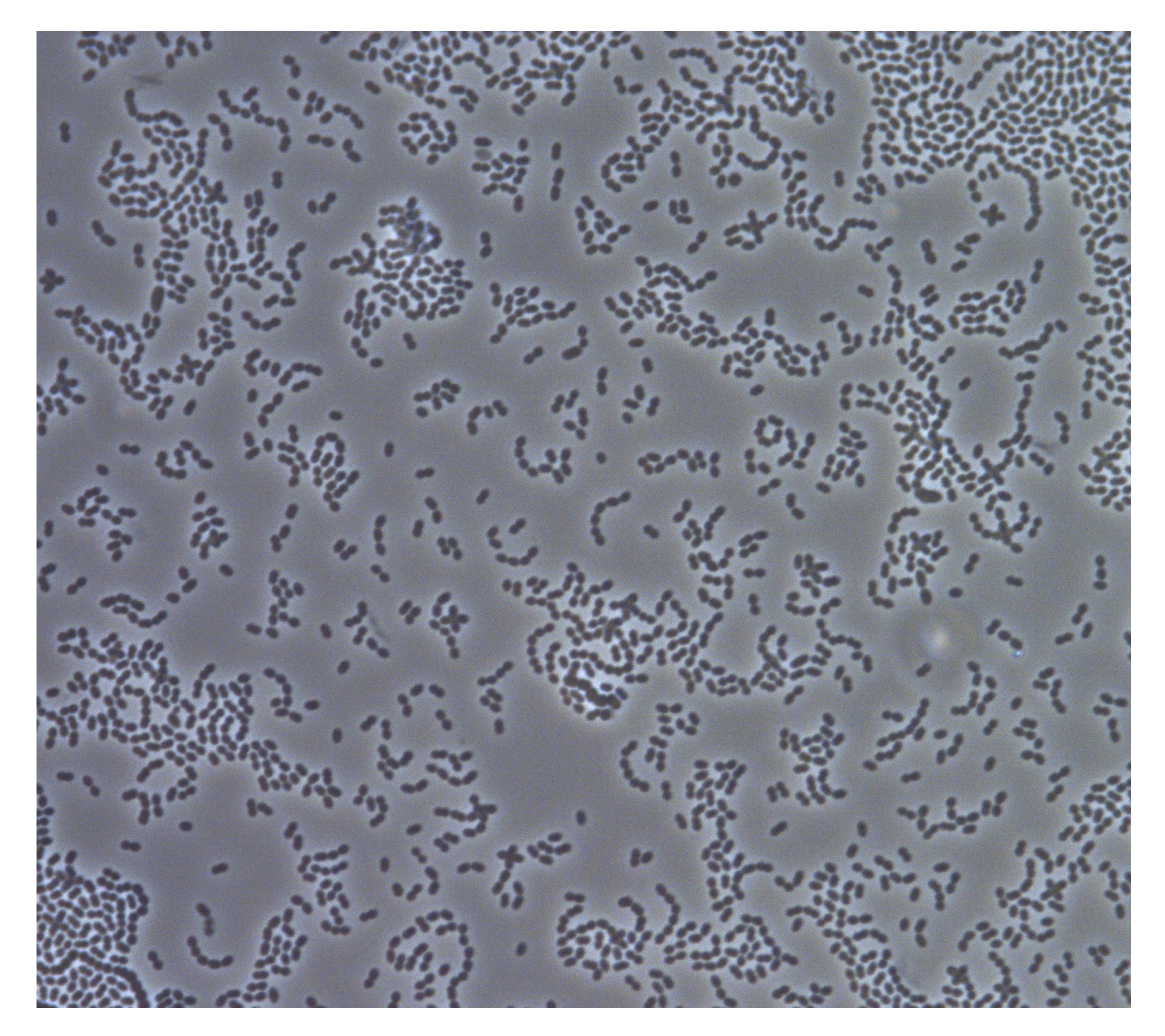

### Sup_Fig_2.tiff

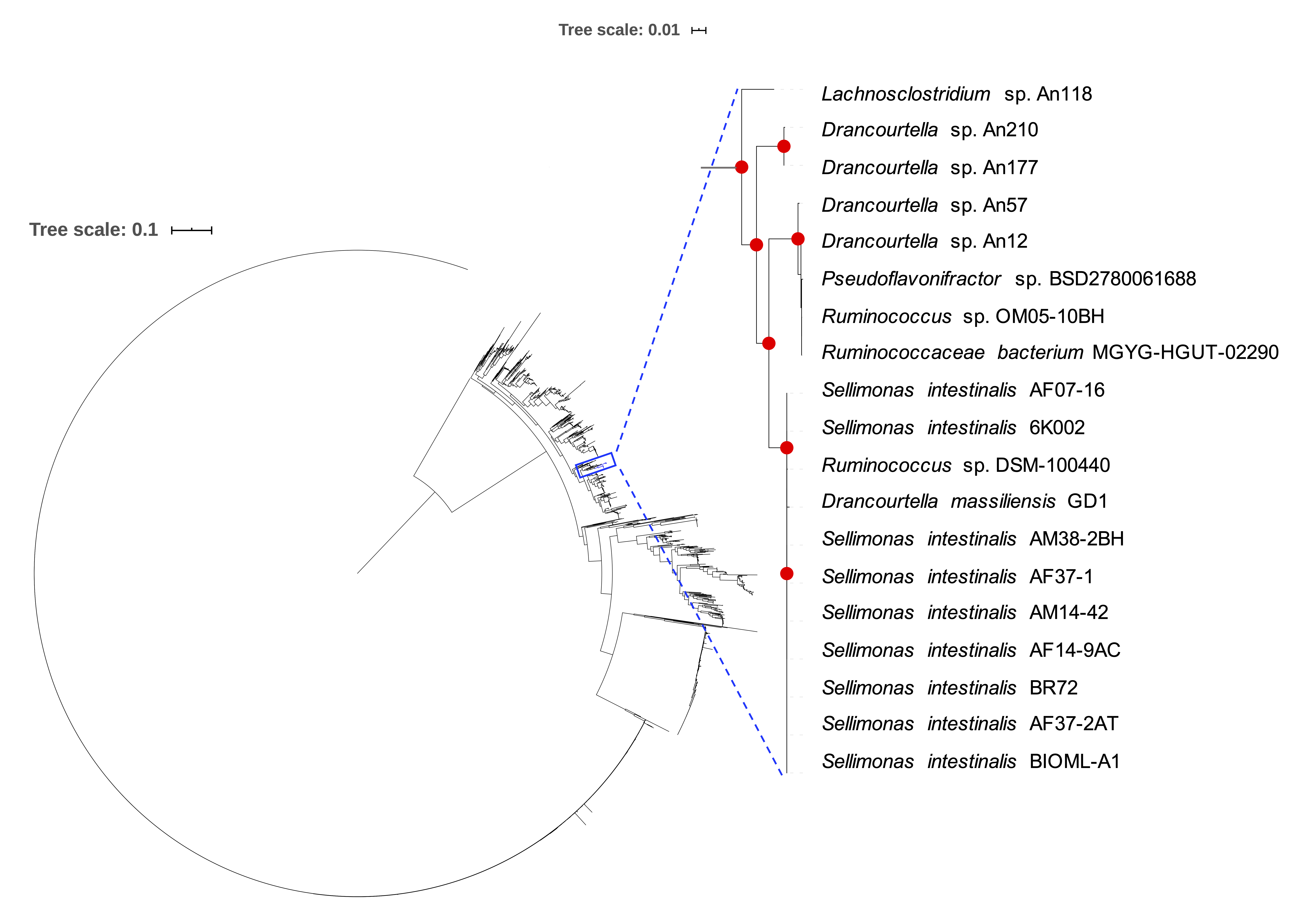

### Sup_Fig_3.tiff

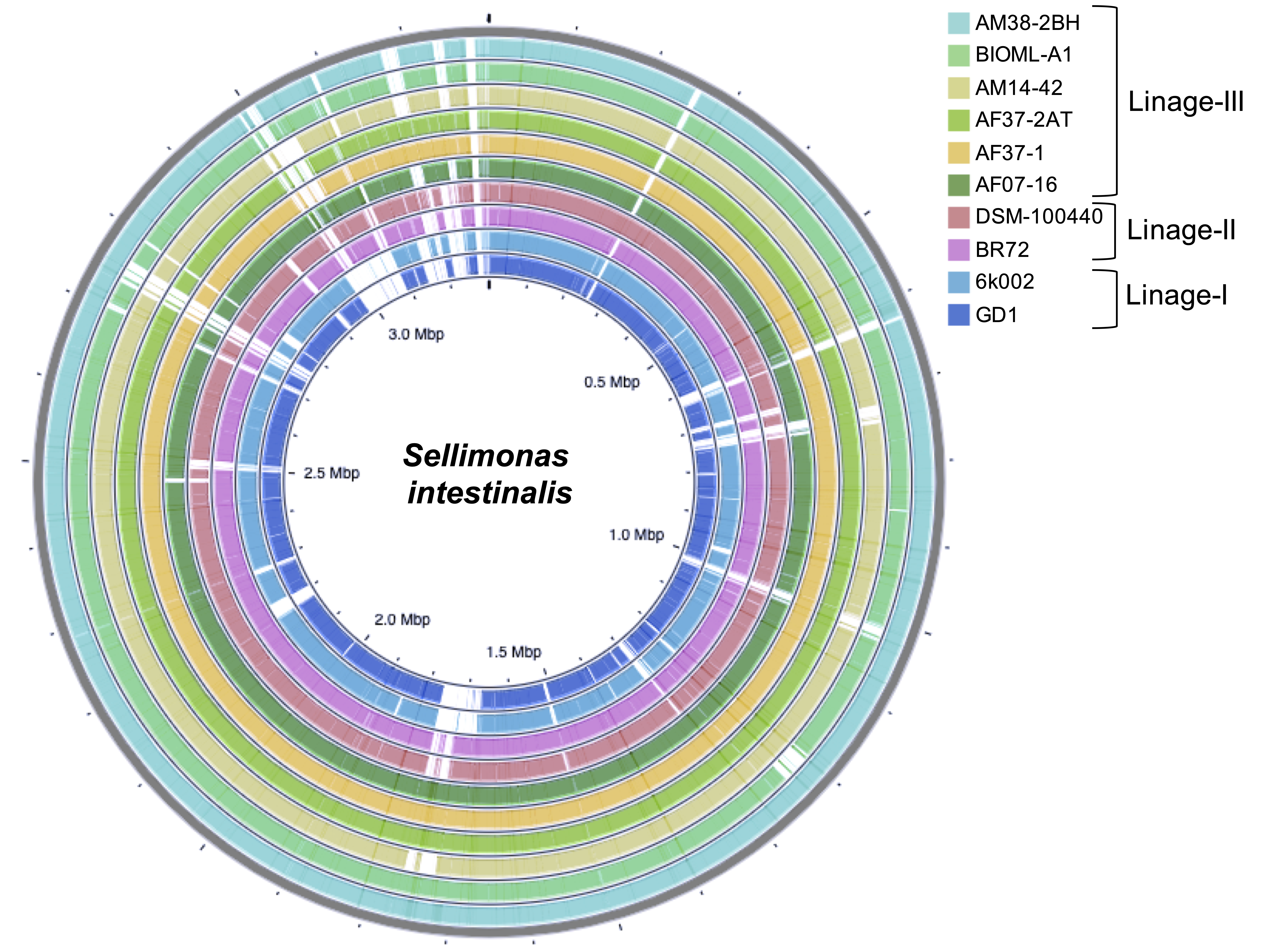
